# Murine metabolic HFpEF is associated with mitochondrial substrate inflexibility and S-nitrosylation remodeling

**DOI:** 10.64898/2026.07.11.737886

**Authors:** Huihui Li, Hanyu Cui, Daiyu Wang, Florian Leuschner, Joerg Heineke, Jiong Hu, Sofia-Iris Bibli

**Affiliations:** Department of Vascular Dysfunction, European Center for Angioscience (ECAS), Medical Faculty Mannheim, Heidelberg University, Mannheim, Germany; Division of Cardiology, Department of Internal Medicine, Tongji Hospital, Tongji Medical College, Huazhong University of Science and Technology, Wuhan, China; Department of Histology and Embryology, School of Basic Medicine, Tongji Medical College, Huazhong University of Science and Technology, Wuhan, China; Department of Cardiology, Internal Medicine III, Heidelberg University Hospital, Heidelberg, Germany; German Center of Cardiovascular Research (DZHK), Partner site Heidelberg-Mannheim, Germany; Department of Cardiovascular Physiology, European Center for Angioscience (ECAS), Medical Faculty Mannheim, Heidelberg University, Mannheim, Germany; Helmholtz-Institute for Translational AngioCardioScience (HI-TAC) of the Max Delbrück Center for Molecular Medicine in the Helmholtz Association (MDC) at Heidelberg University, Heidelberg 69117, Germany

**Author notes:** Correspondence: Huihui Li, MD, PhD. Department of Vascular Dysfunction, European Center for Angioscience (ECAS), Medical Faculty Mannheim, Heidelberg University, Mannheim., Sofia-Iris Bibli, PhD. Department of Vascular Dysfunction, European Center for Angioscience (ECAS), Medical Faculty Mannheim, Heidelberg University, Mannheim, Ludolf-Krehl-Straße 13-17, 68167. These authors contributed equally (Huihui Li, Hanyu Cui).

## Abstract

Heart failure with preserved ejection fraction (HFpEF) is a heterogeneous condition with incompletely defined myocardial mechanisms. Here, using a two-hit murine model of cardiometabolic HFpEF induced by high-fat diet and endothelial nitric oxide synthase inhibition, we define a mitochondrial metabolic phenotype characterized by substrate inflexibility, redox stress, and S-nitrosylation remodeling. While global proteomic changes were modest, metabolomic profiling revealed accumulation of tricarboxylic acid cycle intermediates, increased dicarboxylic acids, and altered redox-associated metabolites, consistent with inefficient oxidative metabolism and mitochondrial redox imbalance in this experimental setting. S-nitrosylation proteomics demonstrated a highly organized and bidirectional remodeling pattern affecting proteins involved in fatty acid/lipid metabolism, carbohydrate metabolism, mitochondrial energy metabolism, amino acid and organic acid metabolism, nucleotide/cofactor metabolism, and redox defense. Beta-hydroxybutyrate (BHB), an alternative mitochondrial substrate, improved basal and ATP-linked respiration, reduced selected TCA-cycle intermediates, lowered mitochondrial reactive oxygen species and the NADH/NAD^+^ ratio, partially restored the GSH/GSSG ratio, and improved diastolic and functional phenotypes without altering preserved ejection fraction. Together, these findings define a redox-sensitive mitochondrial metabolic state in the HFD/L-NAME model and identify ketone supplementation as a partial metabolic rescue strategy in this context. At the same time, they highlight an important limitation of murine HFpEF models: such models do not faithfully reproduce the metabolic phenotype of human HFpEF and should therefore be interpreted as experimental systems rather than human disease equivalents.

## 1. Introduction

Heart failure with preserved ejection fraction (HFpEF) remains mechanistically elusive and therapeutically challenging [1, 2]. Although metabolic stress is a major contributor to HFpEF in a substantial proportion of patients, the mechanisms linking obesity, insulin resistance, inflammation, and nitric oxide dysregulation to impaired cardiac relaxation remain incompletely elucidated [2]. To investigate these mechanisms, several experimental models have been developed [3], among which the high-fat diet/L-NAME two-hit model is widely used because it recapitulates not only preserved ejection fraction, diastolic dysfunction and reduced exercise capacity, but also the systemic metabolic and inflammatory milieu, including obesity, glucose intolerance, hypertension, and nitric oxide dysregulation, that characterizes clinical cardiometabolic HFpEF endotype [4].

Mitochondrial dysfunction has emerged as a recurring feature in HFpEF, but the basis of this dysfunction remains unclear [5]. Emerging evidence suggests that regulatory events at the level of protein function may contribute importantly to the pathophysiology of HFpEF [6, 7]. In this disease context, cardiac mitochondrial proteins are susceptible to acute post-translational modifications (PTMs) and redox-dependent regulatory changes. These processes can interact bidirectionally with reactive oxygen species (ROS), thereby disrupting bioenergetic efficiency and amplifying mitochondrial stress [8]. S-nitrosylation is of particular interest in this context because it can directly modulate enzymatic activity, substrate utilization, and redox buffering [9]. We therefore combined global proteomics, S-nitrosylation proteomics, and metabolomics in a two-hit murine model of cardiometabolic HFpEF. Using this approach, we define a model-specific phenotype characterized by limited proteomic remodeling but pronounced metabolic and S-nitrosylation changes affecting mitochondrial and metabolic pathways. We further show that altering mitochondrial substrate utilization partially improves mitochondrial and functional readouts in this context.

Together, these findings support the concept that the HFD/L-NAME model engages a redox-sensitive mitochondrial metabolic state and provide evidence for interpreting this widely used murine system as a tool to study cardiometabolic stress responses, while recognizing that it does not necessarily reproduce the metabolic organization of human HFpEF.

## 2. Methods and Data Availability

### 2.1 Data availability

The mass spectrometry proteomics data have been deposited to the ProteomeXchange Consortium via the PRIDE partner repository with the dataset identifier PXD075362.

### 2.2 Animal studies

Metabolic HFpEF was induced and functionally characterized as described previously [4]. At the end of experiments, the mice were euthanized, the heart was removed and put into liquid nitrogen immediately. Afterwards, heart tissue was pulverized in liquid nitrogen and separated into three parts for global proteomics, metabolomics and S-nitrosylation proteomics. BHB-Na was added in the drinking water since the first day of diet initiation: 2% w/v ≈ 80–120 mg/day intake. Transthoracic echocardiography was performed with ultrasonography using a Vevo3100 system (Visualsonics, Toronto, Canada) as previously described [10]. Cardiovascular parameters were analyzed using the Vevo LAB desktop software. General exhaustion monitoring was assessed as previously described [11]. In brief, mice were acclimated to a rodent treadmill for three days prior to testing. A progressive maximal exercise test was performed in which treadmill speed was gradually increased until a maximum of 18 m/min was reached. Mice ran until exhaustion, defined as remaining on the shock grid for >5 s without resuming running. Total running time and distance were recorded.

### 2.3 Proteomics and S-nitrosylation modification proteomics

For global proteomics, tryptic peptides were desalted and analyzed by data-independent acquisition (DIA) LC–MS/MS on an Orbitrap Astral mass spectrometer coupled to a Vanquish Neo UHPLC system (Thermo Fisher Scientific).

For S-nitrosylation proteomics, heart tissues were lysed in SDS-containing buffer, and free thiols were blocked with iodoacetamide. S-nitrosylated cysteine residues were selectively reduced with sodium ascorbate and derivatized with a Cys-Tag (CPT) reagent. After tryptic digestion, CPT-labeled peptides were enriched using immobilized metal affinity chromatography (IMAC) and analyzed by DIA LC–MS/MS on the same Orbitrap Astral platform. Raw data were processed using Spectronaut, and S-nitrosylation sites were quantified by intensity-based label-free quantification. PTM sites with at least three valid quantitative values in either group were retained. Differential expression analysis was performed using the limma package in R (|log2 fold change| > 1, P < 0.05), and functional enrichment analysis was conducted using the clusterProfiler R package.

### 2.4 Metabolomics

Metabolites were extracted and analyzed by LC-MS/MS as previously described [12]. Metabolites with QC sample RSD >30% were removed. Differential expression analysis was performed using the limma package in R (|log2 fold change| > 1, P < 0.05). Functional enrichment analysis was performed using MetaboAnalyst.

### 2.5 Cardiomyocyte isolation

Cardiomyocytes were isolated from mice using a modified Langendorff-free method for further evaluation as previously described [13, 14].

### 2.6 Mitochondrial respiration assay

The oxygen consumption rate (OCR) was measured as previously described [13].

### 2.7 MitoSOX measurement

Cells were incubated with MitoSOX™ Red reagent (1:5000 in PBS with Ca^2+^/Mg^2+^) for 30 min at 37°C, and fluorescence was quantified by flow cytometry analysis (BD FACSCanto™ II). Viable single cells were gated based on FSC/SSC characteristics. Data were analyzed using FlowJo V10.

### 2.8 NADH/NAD^+^assay

The assay was performed in isolated cardiomyocytes with a commercially available NADH/NAD^+^ assay kit from Abcam (Berlin, Germany, Cat.No. ab176723) according to the manufacturer’s protocol as described previously [13].

### 2.9 Statistical analysis

All data are presented as the mean ± SD. One-way analysis of variance (ANOVA) followed by Tukey’s multiple comparisons test was used. Details are presented on each figure legend. All analyses were performed using GraphPad PRISM data analysis software (version 10.2; GraphPad Software).

## 3. Results

### 3.1 Cardiometabolic HFpEF shows modest proteomic remodeling but a metabolic signature consistent with mitochondrial substrate inflexibility

Using the two-hit HFD/L-NAME model of metabolic HFpEF, we found modest global proteomic alterations (Fig. 1A, Table S1). Differentially expressed proteins were enriched in cytoskeletal/contractile components (MYH11, TAGLN, CNN1, VIM) and extracellular matrix (COL5A1, COL6A1, COL6A6, COL14A1) (Fig. 1B, C, Table S1), indicating coordinated structural remodeling despite limited abundance-level proteome reprogramming. In contrast, metabolomic profiling revealed a more prominent shift in pathways linked to mitochondrial substrate handling (Fig. 1D–E, Table S2). Tricarboxylic acid (TCA) cycle intermediates were significantly increased, indicating accumulation of mitochondrial carbon intermediates. Concurrently, dicarboxylic acids (azelaic acid, suberic acid) were elevated, suggesting enhanced fatty acid flux and ω-oxidation overflow, consistent with lipid overload. Redox-associated metabolites, including reduced glutathione, were also increased, reflecting activation of compensatory antioxidant mechanisms. In contrast, multiple ω-3 polyunsaturated fatty acid–derived oxylipins were decreased, pointing to impaired protective lipid signaling and altered fatty acid handling (Fig. 1E, F). However, in human HFpEF, some TCA-cycle intermediates such as succinate and fumarate are reduced. Likewise, fatty acid-related metabolites such as medium- and long-chain acylcarnitines, as well as fatty acid oxidation proteins and genes, are generally decreased in human HFpEF [15]. Therefore, the murine and human findings may represent distinct manifestations of impaired mitochondrial fuel flexibility in HFpEF. The murine model reflects a state of substrate overload with redox-constrained oxidative flux, whereas human HFpEF myocardium may represent a more advanced or differently organized state characterized by reduced oxidative organization and impaired substrate utilization capacity.

**Figure 1.**
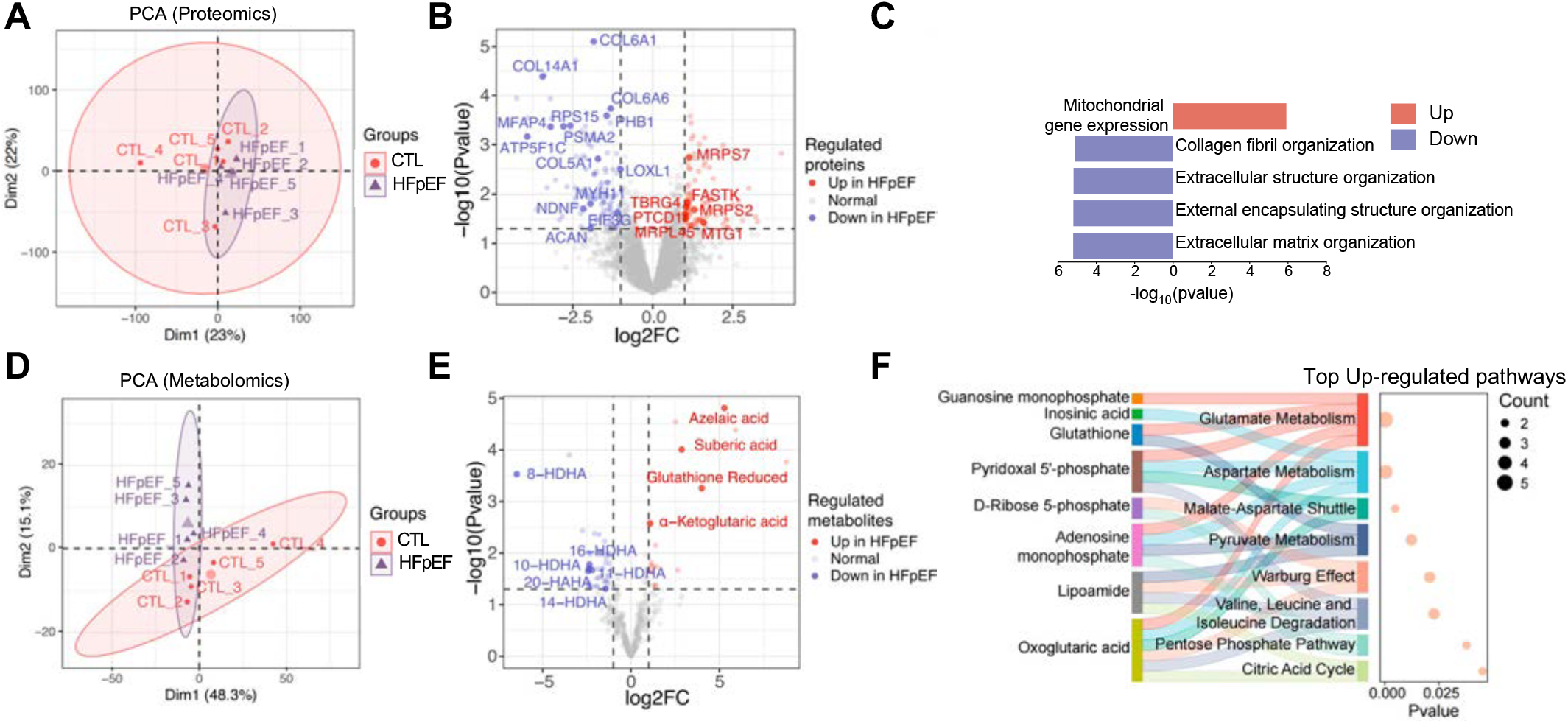
Metabolic HFpEF shows modest proteomic remodeling but marked metabolic evidence of mitochondrial substrate inflexibility. **A–B**. Principal component analysis (PCA) (A) and volcano plot (B) of cardiac proteomes from mice under normal diet (CTL) or treated with high-fat diet and L-NAME (HFpEF). **C**. Bar plot displaying biological pathways enriched among differential proteins between CTL and HFpEF groups. **D–E**. PCA (D) and volcano plot (E) of cardiac metabolomes from the CTL and HFpEF groups. **F**. Integrated bubble plot and Sankey diagram depicting metabolic pathways enriched in HFpEF. Metabolites driving pathway enrichment are highlighted on the left.

Taken together, these data identify a metabolite pattern in the HFD/L-NAME model that is consistent with inefficient oxidative metabolism and mitochondrial redox imbalance. Although steady-state metabolite levels do not allow direct inference of pathway flux, the combined accumulation of TCA-cycle intermediates, altered lipid-associated metabolites, and redox-related changes supports the presence of mitochondrial substrate inflexibility in this experimental setting.

### 3.2 S-nitrosylation remodeling identifies redox-sensitive metabolic control nodes in the HFD/L-NAME model

To explore how metabolic remodeling might be regulated in this experimental setting, we examined cysteine S-nitrosylation as a candidate redox-sensitive post-translational mechanism. Because nitric oxide-related signaling can modulate protein function, enzymatic activity, and substrate utilization through reversible cysteine modification, we asked whether the HFD/L-NAME model exhibits coordinated S-nitrosylation changes across metabolic pathways. Indeed, S-nitrosylation remodeling in HFpEF was significantly altered (Fig. 2A, Table S3). Sites with increased S-nitrosylation were associated with proteins involved in mitochondrial redox defenses and metabolic control, including enzymes regulating fatty acid utilization and oxidative flux such as PDK4, MLYCD, CPT2, and OGDH, as well as contractile and Ca^2+^ handling proteins including Ryr2 and SERCA2 (Fig. 2A, Table S3). Conversely, sites with decreased S-nitrosylation affected complementary modules of contractile structure and fuel handling, consistent with dynamic, site-specific redox regulation of mitochondrial substrate inflexibility (Fig. 2A). Consistent with this, Gene Ontology analysis of the differentially S-nitrosylated proteins identified a strong enrichment for pathways governing catabolic metabolism and energy turnover, including small-molecule catabolism, fatty acid metabolism and oxidation, amino-acid metabolism, organic/carboxylic-acid catabolism, and energy derivation by oxidation of organic compounds (Fig. 2B). Additional enrichment in muscle cell differentiation/development suggests that this metabolic rewiring occurs in parallel with remodeling of cardiac structural programs. Notably, these enrichments were centered on cardiac metabolic processes, rather than broad transcriptional pathways, supporting the concept that HFpEF is characterized by a post-translationally regulated metabolic phenotype. Next, we mapped the differential S-nitrosylation sites of key metabolic proteins across major metabolic modules (Fig. 2C). Distinct and coordinated remodeling was evident in fatty acid/lipid metabolism, carbohydrate metabolism, mitochondrial energy metabolism/TCA cycle, nucleotide and cofactor metabolism, amino acid/organic acid metabolism, and redox/antioxidant/detox pathways. Across these modules, S-nitrosylation changes indicated selective remodeling of metabolic control nodes instead of uniform activation or suppression of entire pathways. Together with the metabolomic alterations described above, these findings are consistent with redox-sensitive post-translational regulation contributing to altered mitochondrial substrate handling in the HFD/L-NAME model.

**Figure 2.**
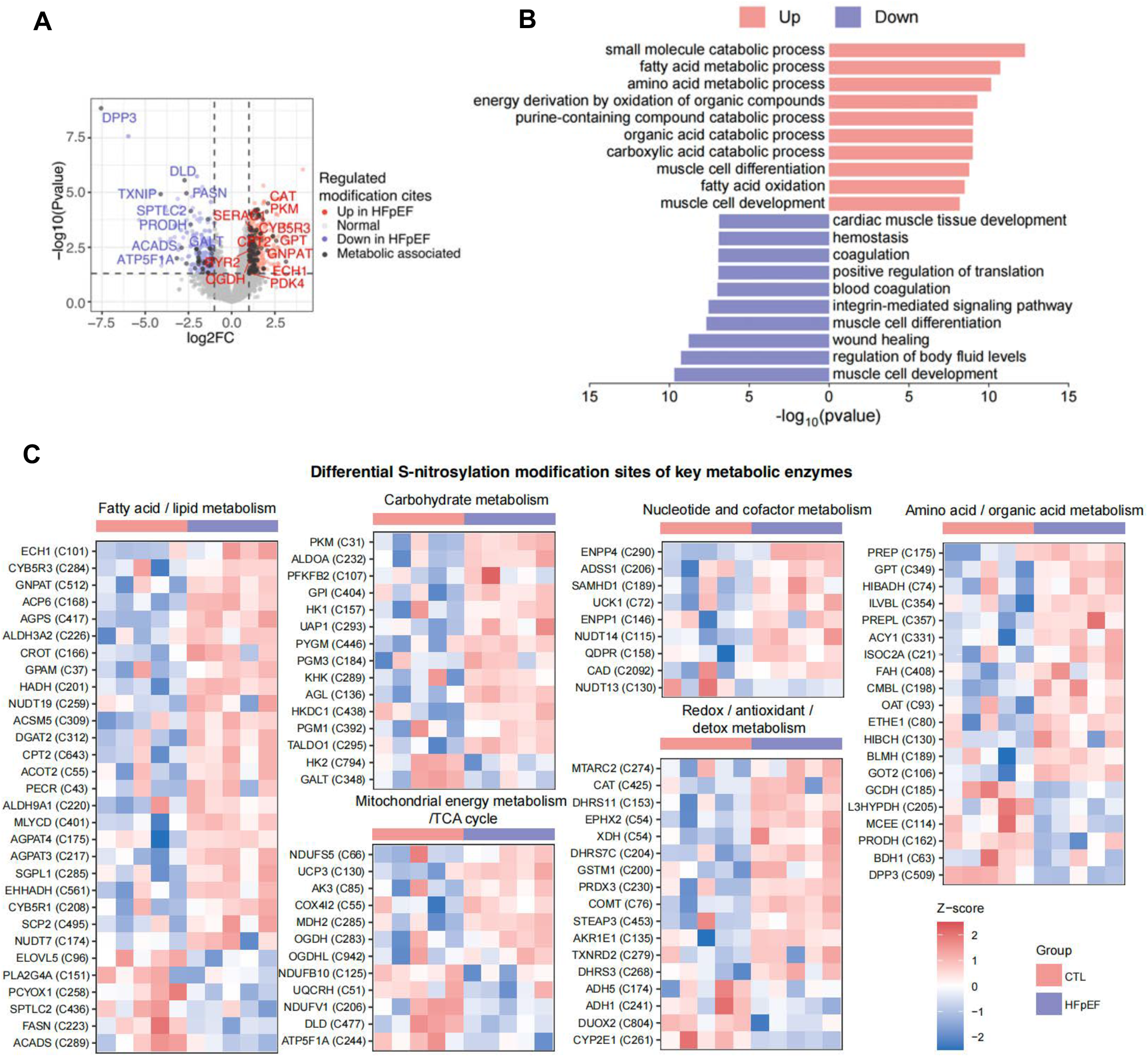
S-nitrosylation is remodeled in HFpEF. **A**. Volcano plot identifying differentially S-nitrosylated cysteines (SNO) in HFpEF compared to CTL hearts. **B**. Bar plot displaying biological pathways enriched among proteins harboring differentially S-nitrosylated cysteines between CTL and HFpEF groups. **C**. Heatmap displaying differentially S-nitrosylated cysteines of key metabolic proteins across major metabolic pathways.

### 3.3 Beta-hydroxybutyrate relieves redox-constrained oxidative metabolism and improves HFpEF phenotypes

To test whether the redox-metabolic state identified in this model could be functionally modified, we examined the effects of shifting substrate availability toward ketone metabolism. Because beta-hydroxybutyrate (BHB) can be converted directly into acetyl-CoA and enter the TCA cycle independently of carnitine-dependent long-chain fatty acid transport [16], we reasoned that ketone utilization might reduce fatty acid-associated redox burden and improve oxidative efficiency. Consistent with this idea, BHB partially restored mitochondrial respiration in HFpEF hearts, as reflected by improved basal and ATP-linked oxygen consumption rates (Fig. 3A). This was accompanied by normalization of TCA-cycle intermediate accumulation, with reductions in aconitate, fumarate, malate, and oxaloacetate, consistent with relief of mitochondrial backpressure and more efficient oxidative flux (Fig. 3B). In parallel, BHB attenuated redox stress, lowering MitoSOX and NADH/NAD^+^, while partially restoring the GSH/GSSG ratio (Fig. 3C–E). Functionally, these metabolic effects translated into improved cardiac performance characterized by better global longitudinal strain (GLS) and lower E/E′, together with reduced heart weight/tibia length and improved exercise capacity, whereas LVEF remained preserved and unchanged, consistent with the HFpEF phenotype (Fig. 3F).

**Figure 3.**
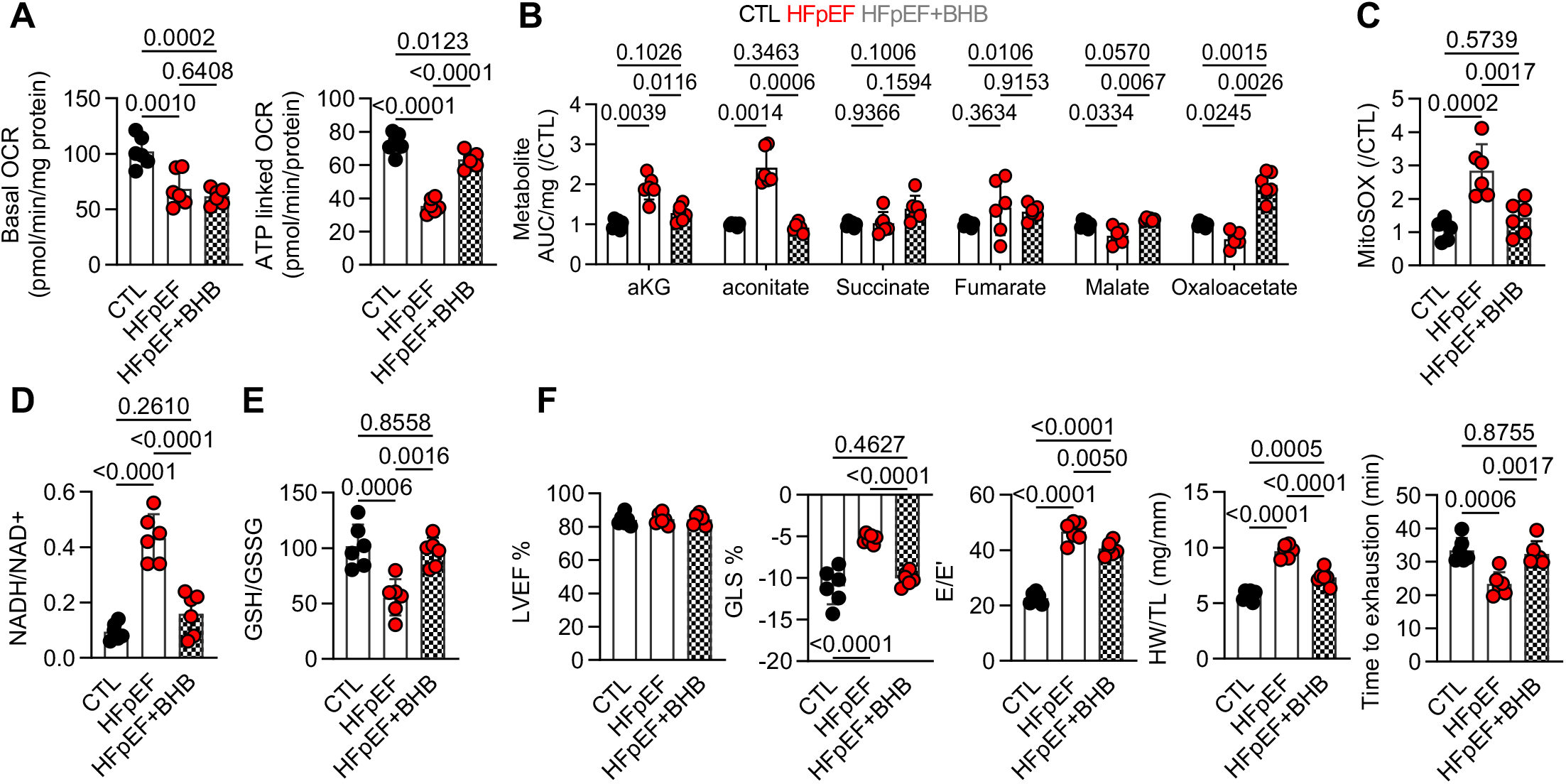
Beta-hydroxybutyrate relieves redox-constrained metabolism and improves HFpEF phenotypes. **A**. Oxygen consumption rate (OCR) from cardiac tissue of CTL and HFpEF mice with or without the addition of beta-hydroxybutyrate (BHB). **B–E**. TCA cycle metabolites (B), MitoSOX (C), NADH/NAD^+^ (D) and GSH/GSSG (E) in isolated cardiomyocytes from CTL and HFpEF mice with or without the addition of BHB. **F**. Percent left ventricular ejection fraction (LVEF%), left ventricular global longitudinal strain (GLS), ratio between mitral E wave and E’ wave (E/E’), ratio between heart weight and tibia length (HW/TL), running time to exhaustion in CTL mice or mice exposed to metabolic HFpEF diet with and without the addition of BHB. Results are presented as mean ± SD. One-way ANOVA, Tukey’s multiple-comparisons test (A–F). n=6/group.

Together, these findings indicate that the redox-metabolic phenotype observed in the HFD/L-NAME model is at least partially reversible. Although these data do not establish the precise metabolic bottleneck engaged by BHB, they support the concept that altering mitochondrial substrate use can partially rescue mitochondrial and functional abnormalities in this experimental setting.

## 4. Discussion

In this study, we define a redox-sensitive mitochondrial metabolic state in the HFD/L-NAME murine model of metabolic HFpEF. Although global changes in the cardiac proteome were relatively modest, integrated metabolomic and nitrosoproteomic profiling revealed mitochondrial substrate inflexibility, redox stress, and structured S-nitrosylation remodeling of key control nodes. Within this experimental context, these data support the interpretation that mitochondrial dysfunction is associated less with broad loss of mitochondrial proteins and more with regulatory remodeling at the level of protein modification and metabolic organization.

Several findings support this interpretation. First, the steady-state metabolomic profile showed accumulation of TCA-cycle intermediates, increased dicarboxylic acids, and altered redox-associated metabolites, together indicating constrained oxidative flux in the setting of persistent substrate oversupply. Although such measurements do not establish pathway directionality or flux, the overall pattern is consistent with inefficient oxidative metabolism and impaired substrate flexibility in this model. This interpretation is further supported by the reduction in mitochondrial respiratory function observed in the Seahorse analysis. Together, these findings suggest that, in the HFD/L-NAME setting, cardiometabolic stress is associated with a mitochondrial state marked by maladaptive substrate handling and redox disequilibrium. Second, our S-nitrosylation proteomics indicate that this metabolic state is accompanied by highly structured and bidirectional redox-sensitive remodeling, rather than a nonspecific increase in oxidative damage. Differential S-nitrosylation was enriched in multiple metabolic modules, including fatty acid/lipid metabolism, carbohydrate metabolism, mitochondrial energy metabolism, amino acid/organic acid metabolism, nucleotide/cofactor metabolism, and redox defense pathways. In parallel, S-nitrosylation changes were also present in proteins involved in contractile and Ca^2+^ handling pathways. This organization is notable because it suggests that metabolic and mechanical remodeling may be linked through shared redox-sensitive regulatory processes. At the same time, our data do not establish the functional directionality of individual S-nitrosylation events, and site-specific validation will be required to determine which modified proteins act as causal control nodes in this model.

An important implication of the present study is that the HFD/L-NAME system should be interpreted as an experimental platform rather than a faithful metabolic surrogate of human HFpEF. Although it is among the most widely accepted and commonly used animal models of HFpEF, it captures only selected aspects of the human HFpEF, which is highly heterogeneous and exhibits considerable diversity across populations shaped by comorbidity burden, age, sex, and disease duration in ways that are difficult to reproduce in mice [17]. Comparison with human myocardial studies suggests that its metabolic organization may differ in important respects from that observed in human HFpEF. In our study, accumulation of TCA-cycle intermediates together with increased dicarboxylic acids suggests excessive fatty acid-derived substrate pressure with downstream oxidative constraint in HFpEF mice. By contrast, Hahn et al. reported reduced medium- and long-chain acylcarnitines, increased pyruvate, lower succinate/fumarate, and accumulation of BCAAs with reduced downstream catabolites in human HFpEF myocardium, a pattern more consistent with impaired substrate utilization capacity and reduced oxidative organization [15]. Thus, while both datasets indicate mitochondrial inflexibility, the murine model may reflect a state of substrate overload and redox backpressure, whereas human HFpEF may exhibit a broader failure of coordinated fuel utilization. We therefore view the present findings not as evidence that murine HFpEF models recapitulate the human metabolic syndrome, but rather as evidence that this widely used model engages a distinct and experimentally tractable redox-metabolic program under cardiometabolic stress. In this sense, the divergence from human HFpEF is not merely a limitation, it is also informative, because it clarifies what this model can and cannot be used to study.

The use of ketones as a therapeutic strategy in HFpEF has also been reported by other groups, although the mechanistic emphasis and outcomes have differed across studies. The recent study by Sun et al. (2025) showed that unlike in HFrEF, myocardial ketone metabolic remodeling may not be a prominent feature of HFpEF. However, this does not exclude the possibility that exogenous ketone supplementation may exert beneficial effects through acute bioenergetic, hemodynamic, or extracardiac mechanisms in selected HFpEF phenotypes [18]. In HFD/L-NAME model, Liao et al. showed that BHB improved diastolic dysfunction and cardiac remodeling, and interpreted these effects mainly in the context of increased cardiac Treg cells and suppression of the NOX2/GSK-3β pathway, thus emphasizing an anti-inflammatory and immunomodulatory mechanism [19]. Deng et al. likewise reported that increasing BHB availability ameliorated HFpEF, and linked this benefit primarily to suppression of a mitochondrial hyperacetylation-NLRP3 inflammasome circuit [20]. Our study differs from these prior reports in both question and mechanistic framework. Rather than focusing primarily on inflammation or immune regulation, we examined BHB in the context of redox-sensitive post-translational control of mitochondrial metabolism. In our study, BHB was used because it offered a direct way to test our central hypothesis that HFpEF is characterized by a redox-constrained oxidative flux that may be relieved by shifting substrate entry into mitochondria. Within this framework, the effects of BHB in our study are mechanistically informative. BHB improved basal and ATP-linked respiration, reduced accumulation of selected TCA-cycle intermediates, lowered mitochondrial ROS and the NADH/NAD^+^ ratio, partially restored the GSH/GSSG ratio, and improved diastolic and functional phenotypes. These changes suggest that BHB did not simply act as a generic alternative fuel, but rather alleviated a state of redox-constrained mitochondrial substrate inflexibility. The differences among studies likely reflect differences in HFpEF models, treatment modality, exposure duration, and the specific metabolic defect being interrogated. Importantly, our findings do not contradict prior work. Instead, they place ketone therapy into a broader mechanistic context in which anti-inflammatory signaling, mitochondrial acetylation, and ketone oxidation may all intersect with redox-sensitive control of mitochondrial metabolism.

Our study has several limitations. We focused on a specific cardiometabolic model of HFpEF and therefore cannot assume that all HFpEF phenotypes are driven by identical mechanisms. In addition, although the combined steady-state metabolomics, Seahorse analysis, and rescue experiments strongly support impaired oxidative flexibility, we did not directly measure substrate flux in vivo. Likewise, while our data implicate S-nitrosylation as a central regulatory mechanism, site-specific functional validation of individual modified enzymes will still be required to establish direct causal links between particular nitrosylation events and altered metabolic flux.

Despite these limitations, the integrated evidence presented here supports a coherent model in which the HFD/L-NAME model engages a redox-sensitive mitochondrial metabolic state characterized by altered substrate handling, redox imbalance, and structured S-nitrosylation remodeling. More broadly, the study highlights both the usefulness and the limits of current murine HFpEF systems: they can reveal mechanistic responses to cardiometabolic stress, but they should not be assumed to faithfully reproduce the metabolic architecture of human HFpEF. Within that boundary, our findings identify reversible redox-metabolic organization as a feature of this model and support metabolic interventions as a means to probe and partially modify that state.

## Table legends

**Table S1** *Proteomic expression matrix and differential proteins in hearts of CTL versus HFpEF mice*

**Table S2** *Metabolomic matrix and differential metabolites in hearts of CTL versus HFpEF mice*

**Table S3** *S-nitrosylation proteomic matrix and differentially modified sites in hearts of CTL versus HFpEF mice*

## Acknowledgments

The authors are indebted to Britta Heckmann and Natalie Klinger for expert technical assistance.

